# Seesaw protein: Design of a protein that adopts interconvertible alternative functional conformations

**DOI:** 10.1101/2024.05.25.595880

**Authors:** Toma Ikeda, Tatsuya Nojima, Souma Yamamoto, Ryusei Yamada, Hiroki Konno, Hideki Taguchi

## Abstract

Classical Anfinsenʼs dogma states that a protein folds into a single unique conformation with minimal Gibbs energy under physiological conditions. However, recent advances have revealed that single amino acid sequences can fold into two or more conformations. Here, we propose a novel approach to design a protein that adopts interconvertible alternative functional conformations, termed “seesaw” protein (SSP). An SSP was engineered by fusing GFP lacking the C-terminal β-strand and DHFR lacking the N-terminal β-strand with an overlapping linker, which can be competitively incorporated into either the GFP or the DHFR moiety. In vivo and biochemical analysis, including AFM imaging, demonstrated that the SSP adopts two alternative conformations, which can be biased by point mutations and ligand binding. In addition, the balance of the seesaw can be reversibly changed depending on buffer conditions. In summary, our design strategy for SSP provides a new direction for creating artificial proteins with on-off behaviors.

## Introduction

The functional three-dimensional structure of a typical globular protein is established by folding the polypeptide chain, with a naturally selected specific amino acid sequence, into its unique minimum free energy state. This principle of protein structure formation—the one-to-one correspondence between a specific amino acid sequence and a specific structure—is known as Anfinsen’s dogma (1). However, the dogma that the conformation is uniquely determined by the amino acid sequence has been challenged. Protein conformation is not always uniquely defined. Pioneering work, such as the discovery of “switch peptides”, whose secondary structure varies depending on environmental factors like pH or temperature (2, 3), and the “Chameleon” sequence, where the same amino acid sequence forms distinct secondary structures when incorporated into different proteins (4, 5), has demonstrated that a particular amino acid sequence can adopt multiple secondary structures in a context-dependent manner.

In addition to those secondary structural switches, it is now clear that proteins with multiple conformations from single sequences exist in nature. For example, amyloids, which irreversibly change their conformation from a certain conformation to intermolecular β-sheets, are a well-known example (6). Recently, it has also been reported that there are proteins such as Morpheeins, which drastically change their tertiary structure depending on their quaternary structures (7–9), metamorphic proteins(10–12), and fold-switch proteins (13, 14), which can reversibly change their topology depending on environmental factors. In fact, more than 100 conformation-changing proteins have been reported, and about 4% of the proteins in the PDB are assumed to have multiple conformations(15). These findings imply that Anfinsen’s dogma cannot fully explain all aspects of the protein world. These proteins have been reported to undergo allosteric regulation and acquire new functions through conformational changes, suggesting that they may have physiological significance (16–18).

Beyond conformation-changing proteins in nature, the artificial design of proteins with multiple structures, including de novo design proteins (19–22), has advanced to elucidate and harness functional controls through conformational changes (23–29). Using natural proteins as templates, Solomon *et al*. designed a temperature-dependent conformational switch protein by precisely matching the sequence homology of a 56-amino-acid protein with a 3α fold and a 95-amino-acid protein with an α/β-plait topology (30). Sekhon *et al*. designed a protein in which the C-terminus of ribonuclease barnase was duplicated and fused to the N-terminus (31). One of the C-terminal regions is inactivated by mutation, enabling the RNase activity to be controlled in an on/off manner. Sander *et al*. fused the N-terminal regions of two related fluorescent proteins, Venus and Cerulean, with a single C-terminal region that shares a common sequence (32). They succeeded in modulating the yellow-to-blue fluorescence ratio by regulating the translation elongation rate through synonymous substitutions.

Expanding on these strategies for switching conformations, a new approach to control functional switching by merging two proteins with distinct functions would be invaluable in the era of creating de novo design proteins. Consequently, we devised a method to fuse two proteins, GFP and DHFR, by overlapping their C- and N-terminal regions, thereby overlapping a segment of their C- and N-termini, respectively. We named the resulting fusion protein “Seesaw protein (SSP)” because its structure and function can be reciprocally altered by integrating either the C-terminus or the N-terminus of the design protein. The two states of the SSP were found to be adjustable by various factors, including mutagenesis and buffer conditions.

## Results

### The design of a protein capable of adopting two alternative conformations

We designed a protein that adopts alternative functional conformations by developing a fusion method for two proteins, overlapping them at the ends. Specifically, a segment of the C-terminal region of the first protein is overlappingly fused with that of the N-terminal region of the second protein (Fig. 1A). If the overlapping region (OR) can complement both proteins, the resulting design protein is expected to possess two alternative states depending on whether the OR is incorporated into the N-terminal or C-terminal moieties. The length of the OR was adjusted to be sufficient to include essential regions for the native states and functions of both parent proteins. Consequently, the designed protein may exhibit two different functional states depending on the region of the designed protein interacting with the OR. Additionally, we anticipated that mutations altering the stability of each moiety, ligands, and buffer conditions would induce changes in the two conformations (Fig. 1B). As the two different functional states controlled by slight differences reminded us of a seesaw, we named such a fusion protein “seesaw protein (SSP)”.

**Figure 1.**
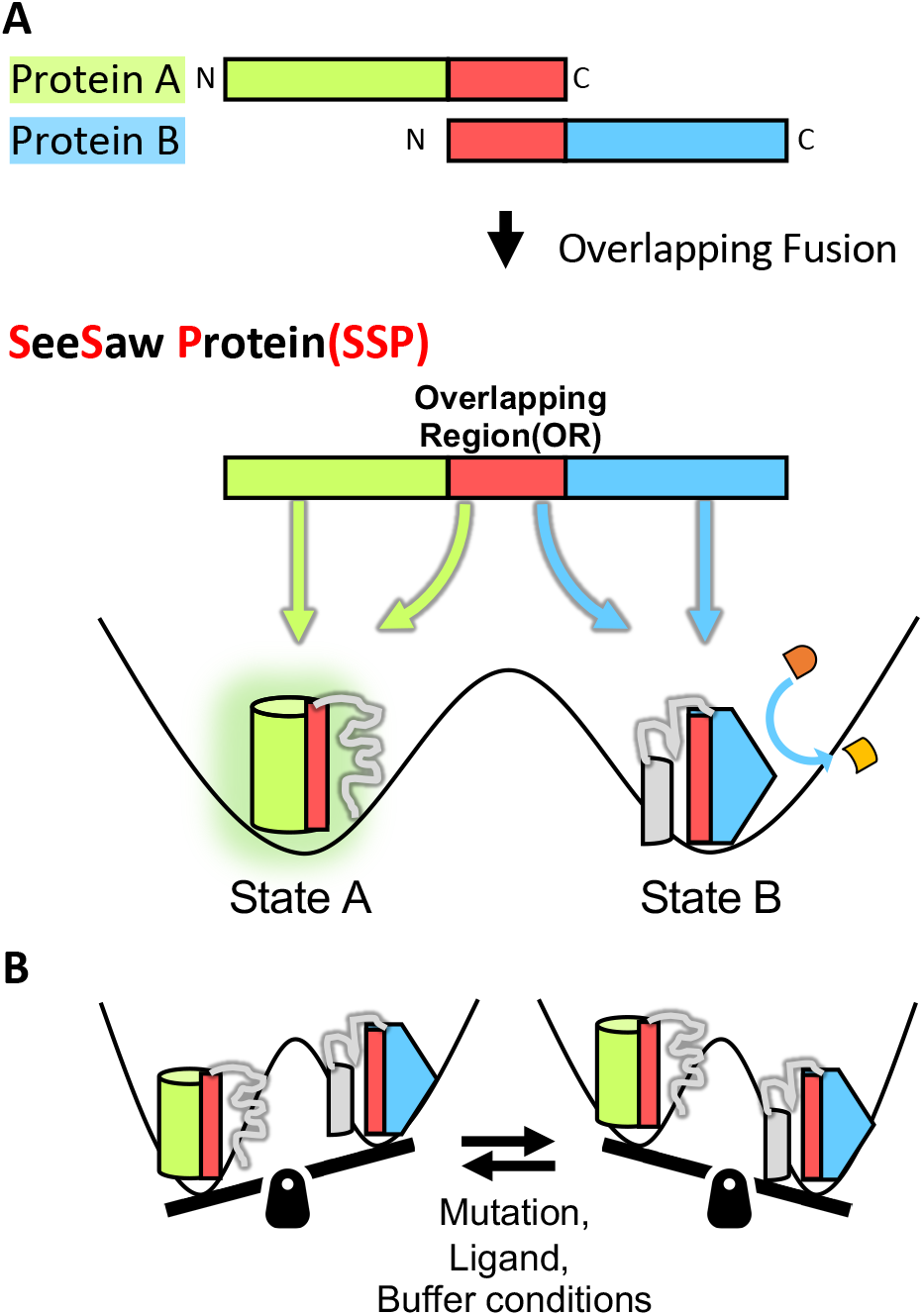
Design principle of a Seesaw Protein (SSP): a protein with interconvertible two alternative functional states. (*A*) Fusion of Two Proteins: Two proteins with different functions, A and B, are joined together by overlapping their ends. In this illustration, the C-terminal region of protein A overlaps with the N-terminal region of protein B and is fused together. If the overlapping region (OR) is crucial for the function of each parent protein, then only the moiety to which the OR binds should exhibit a function. Therefore, two functional states, states A and B, are achieved depending on whether the OR binds to either moiety. This fusion protein is termed a seesaw protein (SSP) because it exhibits seesaw-like behavior, switching between the two functional states within a single amino acid sequence. (*B*) Switchable States of SSP: The SSP can switch between two states. The seesaw tilts toward the more stable state state of the two states of the designed protein. The balance can be adjusted by mutations, ligands, buffer conditions, and other factors.

### Creation of a “seesaw” protein (SSP_GD_) using GFP and DHFR

To create an SSP, we employed superfolder GFP and *Escherichia coli* dihydrofolate reductase (DHFR) as parent proteins (Fig. 2A). These proteins were chosen based on three criteria. First, they were selected for being monomeric proteins to simplify the equilibrium state of each moiety. Second, to facilitate the detection of the two different functional states, proteins with measurable function/activity in vivo were preferred: the GFP state can be evaluated via fluorescence imaging. For DHFR, we devised a colony assay system utilizing a DHFR inhibitor, trimethoprim (TMP). Specifically, under conditions where endogenous DHFR, essential for growth, is inhibited by TMP and therefore unable to promote growth, the growth defect is complemented only when exogenously expressed active DHFR is present (*SI Appendix*, Fig. S1A). Third, proteins with relatively high stability in the wild type, for which mutants with varying stabilities have been extensively studied, were chosen.

**Figure 2.**
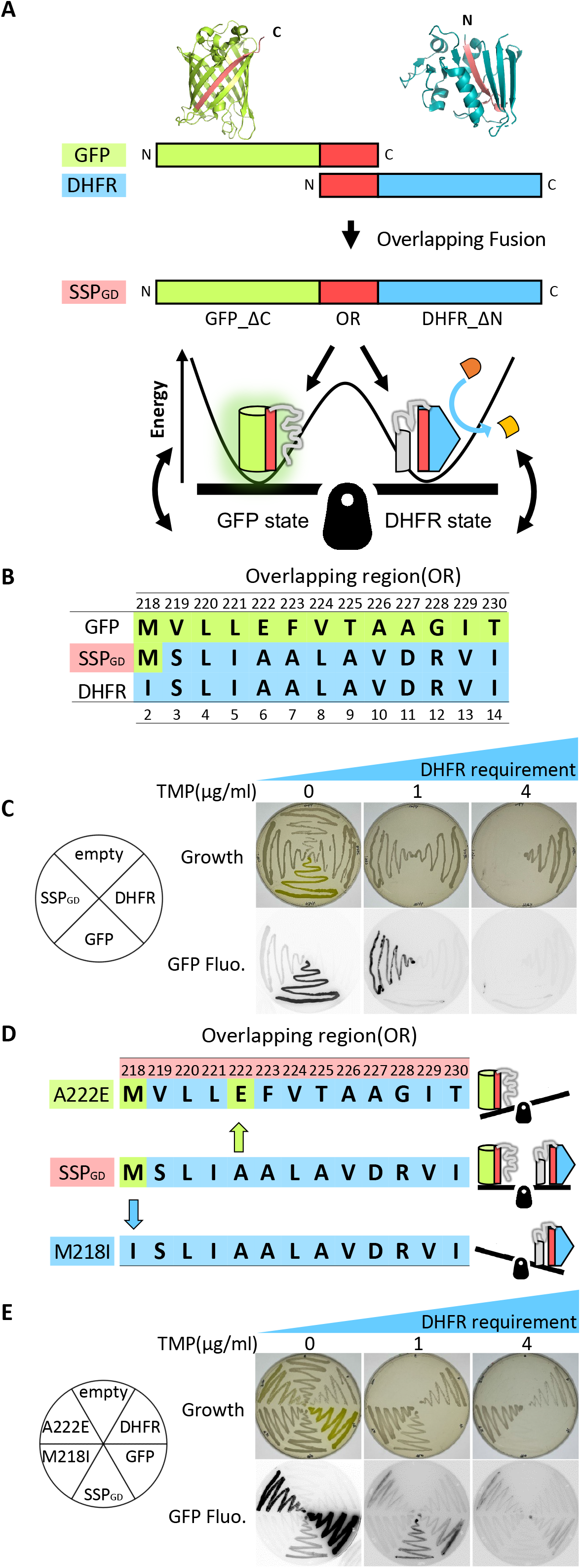
Creation of an SSP (SSP_GD_) composed of GFP and DHFR, exhibiting two distinct functions in the cell. (*A*) Design of the SSP (SSP_GD_) composed of GFP and DHFR. SSP_GD_ consists of three components: GFP lacking a C-terminal region (GFP_ΔC), an overlapping region (OR), and DHFR lacking an N-terminal region (DHFR_ΔN). (*B*) Alignment of the OR with the C-terminal region of GFP and the N-terminal region of DHFR. Amino acids derived from GFP or DHFR are colored green and blue, respectively. The numbers indicate the amino acid residue numbers of GFP and DHFR, with the residue 220th (Leu) common to both. The residue numbers of the SSP_GD_ correspond to those of GFP. (*C*) In vivo colony assay to assess SSP_GD_ functions. The positions of overexpressed proteins in the *E. coli* cells are shown in the circle (*left*). “Empty” refers to cells not overexpressing SSP_GD_ or the parent proteins. After incubation at 37 °C for 6 h, proteins were overexpressed for 72 h at 18 °C in plates with varying TMP concentrations (0, 1, and 4 µg/ml). Plates were photographed (*top*) and imaged for fluorescence (*bottom*). (*D*) Amino acid sequences of ORs in SSP_GD_ and its mutants, SSP_GD222_ and SSP_GD218_, affecting the seesaw balance. The color code is consistent with (*B*). The expected seesaw balances for each SSP protein are illustrated on the right. (*E*) In vivo colony assay to assess SSP_GD222_ and SSP_GD218_ functions. The explanation is the same as described in (*C*).

A GFP-DHFR fusion protein was engineered to include a 13-amino-acid long OR that exclusively shares either the C-terminal strand (218-230) of GFP or the N-terminal strand (2-14) of DHFR (Fig. 2B and *SI Appendix*, Fig. S1B). Initially, we confirmed that monomeric GFP and DHFR lacking the OR (GFPΔC and DHFRΔN) exhibited loss of fluorescence and were not expressed in cells in the presence of TMP, respectively (*SI Appendix*, Fig. S1C). Following a series of chimera analyses (*SI Appendix*, Fig. S1B, D), we found that utilizing the sequence MSLIAALAVDRVI as OR, where the first M is derived from GFP, the third L is common, but the remainder is from DHFR, resulting in both GFP fluorescence and DHFR activity in the colony assay (Fig. 2B, C). We verified that the full-length fusion protein was expressed in a soluble fraction and that degradation, which could produce monomeric DHFR or GFP, was negligible (*SI Appendix*, Fig. S1E), suggesting that the degradation products of the fusion are unlikely to contribute to DHFR resistance. Collectively, these results suggest that the GFP-DHFR fusion protein (SSP_GD_) adopts two alternative functions: the GFP state, in which the GFP region incorporates the OR, and the DHFR state, in which the DHFR region incorporates the OR.

To alter the balance between the two functions in SSP_GD_, mutations were introduced to destabilize one state and stabilize the other (Fig. 2D). To stabilize the DHFR moiety, we substituted Met 218 with Ile (SSP_GD218_), which is the original residue in the corresponding DHFR region (Fig. 2D). For GFP stabilization, the 222^nd^ residue, critical for GFP stability, was replaced with the GFP-derived residue (SSP_GD222_) (Fig. 2D). The results revealed that when the SSP mutants were expressed in *E. coli, E. coli* expressing SSP_GD218_ grew in the presence of higher concentrations of TMP compared to SSP_GD_ and nearly lost GFP fluorescence (Fig. 2E and *SI Appendix*, Fig. S1E). Conversely, *E. coli* expressing SSP_GD222_ failed to grow in the presence of TMP, but exhibited enhanced GFP fluorescence compared to SSP_GD_ (Fig. 2E and *SI Appendix*, Fig. S1E). Thus, single amino acid mutations in the OR appear sufficient to switch between the two different functions.

Subsequently, we examined whether mutations outside the OR could influence the balance of the seesaw. To investigate this, we reverted the mutations introduced for the stabilization in sfGFP, all outside the OR, to wild-type GFP residues in SSP_GD222_, a GFP-biased SSP (33)Out of the six mutations tested, four point mutations exhibited both GFP fluorescence and DHFR activities (*SI Appendix*, Fig. S1F, G), suggesting the existence of multiple pathways for altering the seesaw balance..

### The purified SSP_GD_ exhibits two alternative native conformations

We conducted characterization of the SSPs purified from *E. coli*. Prior to analyzing the SSP fusion proteins, we evaluated the monomeric GFP and DHFR proteins, which replaced the C-or N-terminal regions, respectively, with the OR in SSP_GD_. The DHFR activity of DHFRΔN with OR (OR-DHFRΔN) closely resembled that of the wild-type DHFR monomer (*SI Appendix*, Fig. S2A). In contrast, the fluorescence of GFPΔC with OR (GFPΔC-OR) was only a few percent of that of GFP (*SI Appendix*, Fig. S2A), suggesting that substituting the C-terminal strand with a strand mostly derived from the DHFR sequence significantly compromised the fluorescence property of GFPΔC-OR. Subsequently, we quantified the activity/fluorescence of SSP_GD_, namely GFPΔC-OR-DHFRΔN, in comparison to the monomeric counterparts (Fig. 3A). The DHFR activity of SSP_GD_ was approximately 5% of that of OR-DHFRΔN monomer, whereas the fluorescence of SSR_GD_ was about 85% of that of the GFPΔC-OR monomer. The diminished DHFR activity was not solely attributed to the fusion of DHFR and GFP, as the conventional fusion of GFP and DHFR with a Gly-Ser linker did not significantly reduce the DHFR activity (*SI Appendix*, Fig. S2A), suggesting that the proportion of the DHFR moiety in SSR_GD_ would be low.

**Figure 3.**
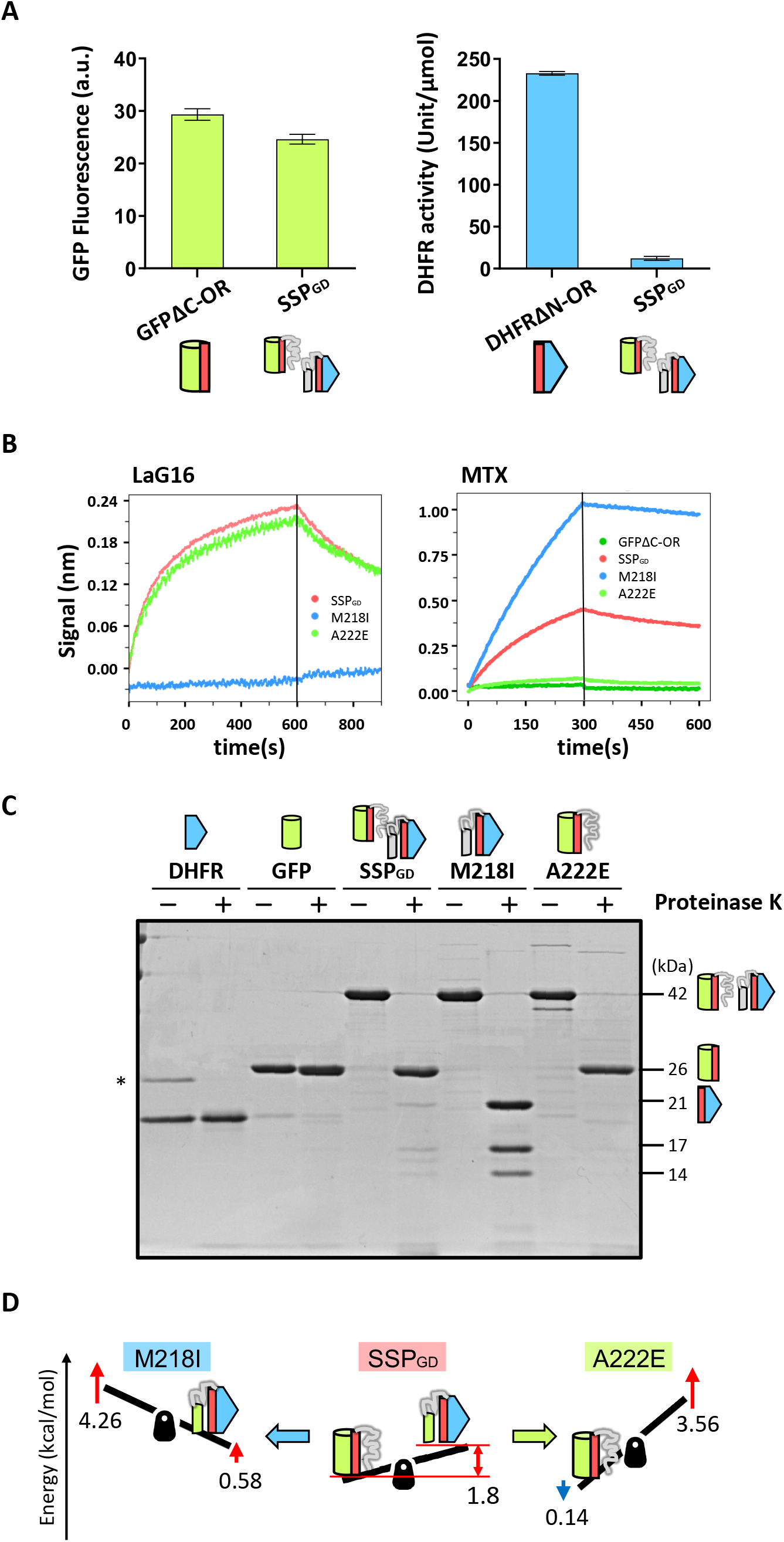
SSP_GD_ exhibited two different conformations. (*A*) Measurement of GFP fluorescence (*left*) and DHFR enzymatic activity (*right*) of SSP_GD_ and its parent protein monomers, depicted schematically as in Fig. 2*A*. Fluorescence excitation and emission wavelength were 480, 513 nm, respectively. Unit is defined as the amount of enzyme degrading 1 µmol of substrate per minute. Data represent the means (±S.D.) of three independent experiments. (*B*) Bio-layer interferometry (BLI) analysis. (*left*) Binding between SSPs and a GFP nanobody (LaG16)-immobilized sensor. The sensor was dipped in a kinetics buffer (PBS supplemented with 0.1% BSA and 0.02% Tween20) containing 50 nM SSPs and then transferred to the kinetics buffer without SSPs. (*right*) Binding between SSPs or GFPΔC-OR and an MTX-labeled sensor. MTX, a DHFR inhibitor, was immobilized onto the SA sensor. The MTX-labeled sensor was immersed in the kinetics buffer containing 2 µM GFPΔC_OR, SSP_GD_, SSP_GD222_, or 20 nM SSP_GD218_, and then transferred to the kinetics buffer without SSPs or GFPΔC_OR. (*C*) Limited proteolysis of SSPs. Purified SSP or mutants (5 µM) were incubated with 1 µg/ml proteinase K at 18 °C for 40 min. After quenching with PMSF, the limited proteolysis products were evaluated by SDS-PAGE. The N-terminal sequences of the limited proteolysis products were determined by Edman degradation (Fig. S2*B*). Schematics on the right side of the gel show the limited proteolysis products corresponding to the bands. The asterisk indicates a contaminant that could not be removed by purification. (D) The thermodynamic balance of the seesaw. The free energy difference between the GFP and DHFR states in SSP_GD_ was analyzed using band intensities of the limited proteolysis products. In addition, changes in thermodynamic stabilities of the two states of SSP mutants relative to SSP_GD_ were predicted by FoldX. The predicted change in thermodynamic stability of each state corresponds to the tilt and height of the seesaw.

Next, we investigated whether SSP_GD_ adopts two alternative conformations using bio-layer interferometry (BLI). We prepared a nanobody (LaG16) that binds to the native GFP structure and immobilized it on a BLI biosensor on an Octet system. The GFP nanobody bound to SSP_GD_ and SSP_GD222_, but not SSP_GD218_, indicating that SSP_GD_ and SSP_GD222_ possessed a native GFP conformation, which was absent in SSP_GD218_ (Fig. 3B *left*). Concerning the DHFR moiety, we used a DHFR inhibitor, methotrexate (MTX), as it is known to specifically bind to the native state of DHFR (34, 35). BLI analysis using a biosensor immobilized with MTX revealed that SSP_GD_ and SSP_GD218_ were bound to the MTX-immobilized sensor (Fig. 3B *right*), indicating that SSP_GD_ and SSP_GD218_ possess native DHFR moiety. In summary, BLI analysis demonstrated that SSP_GD_ and its variants have moieties with native conformations depending on the seesaw balance.

To further confirm the existence of two alternative conformations, we employed a limited proteolysis approach to distinguish between them. Typically, unstructured regions are susceptible to proteases, while stable folded structures remain intact. Under conditions where proteinase K did not degrade monomeric GFP or DHFR, the protease treatment completely digested the full-length SSP_GD_ and its variants (42 kD) (Fig. 3C). The protease-resistant fragments were approximately 26 kD (SSP_GD_ and SSP_GD222_), 21 kD (SSP_GD_ and SSP_GD218_), 17 and 14 kD (SSP_GD218_) (Fig. 3C). Edman degradation analysis and predicted molecular weights identified that the 26 and 21 kD fragments originated from the intact GFP and DHFR moieties, respectively (*SI Appendix*, Fig. S2B). The 17 and 14 kD fragments in SSP_GD218_ included the N-terminal half of GFP (*SI Appendix*, Fig. S2B), indicating that the GFP moiety in SSP_GD218_ possesses a folded structure resistant to proteolysis. Limited proteolysis did not alter the intensity of GFP fluorescence or DHFR activity (*SI Appendix*, Fig. S2C), supporting the degradation of only the unstructured regions in SSP_GD_ and its variants, without affecting GFP fluorescence or DHFR activity. The predominance of GFP- and DHFR-containing fragments in SSP_GD222_ and SSP_GD218_, respectively, suggests an extremely biased seesaw balance in the mutants, consistent with in vivo assay and BLI analysis. Conversely, SSP_GD_ predominantly produced a 26 kD fragment (GFP moiety) and a minor 21 kD fragment (DHFR moiety), indicating that SSP_GD_ adopts two alternative conformations with a strongly biased GFP moiety.

Based on the ratio of the two bands in SSP_GD_ (∼9.5:∼0.5) we calculated the Gibbs energy change (ΔG) for the seesaw slope (DHFR/GFP) to be +1.8 kcal/mol (Fig. 3D). Concerning the mutants, differences in thermodynamic energy between “wild-type” SSP_GD_ and the mutants (ΔΔG) were predicted by FoldX (36, 37). The estimated net ΔG values for SSP_GD218_ and SSP_GD222_ were -1.9 and +5.5 kcal/mol, respectively (Fig. 3D).

### Direct visualization of the two distinct conformations in SSPs by high-speed atomic force microscopy (HS-AFM)

To ascertain that SSPs indeed adopt two distinct conformations, we utilized high-speed atomic force microscopy (HS-AFM) to directly visualize the alternative structures of SSPs. HS-AFM imaging of SSPs immobilized on a mica surface distinctly revealed that SSP_GD222_ displayed a single large domain structure, whereas SSP_GD218_ exhibited a structure with two connected domains -one large and one small (Fig. 4 *left and right*, Movies S1, S2). These structures correspond to an intact DHFR and a protease-resistant segment of GFP, respectively. HS-AFM observations of SSP_GD_ predominantly displayed SSP_GD222_-like structures, occasionally featuring SSP_GD218_-like structures consisting of two domains (Fig. 4 *middle*, Movies S3, S4), consistent with the ratio estimated from the limited proteolysis experiment. Furthermore, the visualization of SSP_GD_ and its variants clearly demonstrated that SSPs can exhibit two distinct states in their monomeric states, thus excluding the possibility of conformational conversion due to oligomer formation. Taken together, we conclude that SSP_GD_ indeed adopts two alternative conformations, which can be biased by single mutations.

**Figure 4.**
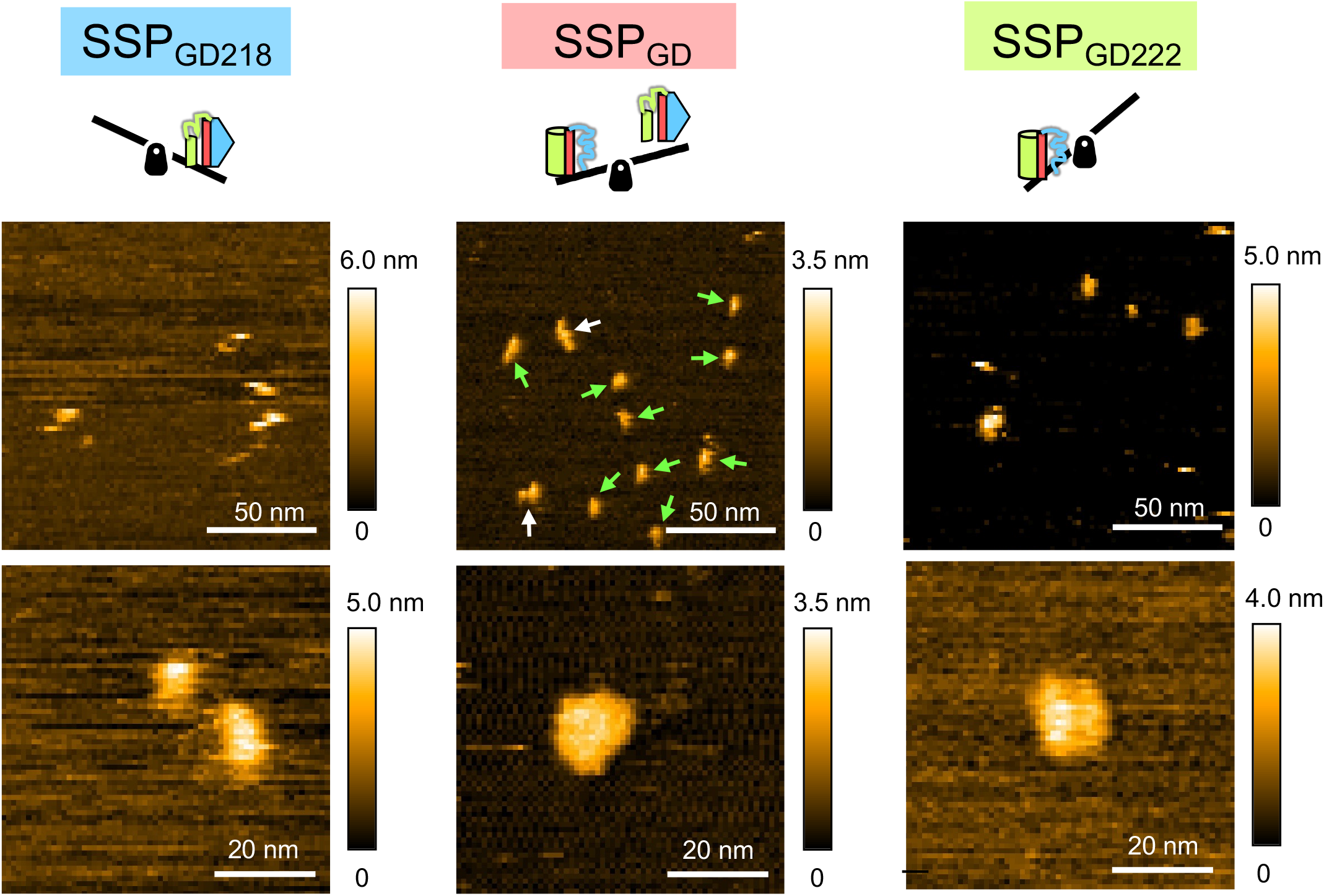
Direct visualization of SSP_GD_ and its mutants using high-speed atomic force microscopy (HS-AFM). HS-AFM imaging of the structural dynamics of SSPs in solution. The top three images show low magnification (wide range) to depict multiple SSPs, while the bottom three present representative magnified images. In the wide range SSP_GD_ image, particles that were considered by visual inspection from the movie to consist of two domains or one domain are indicated by white and green arrowheads, respectively. (*Left*) HS-AFM images of SSP_GD218_. For the top and bottom images, operational parameters were as follows: scanning range, 200 × 200 nm (100 × 100 pixels) and 70 × 700 nm (70 × 70 pixels); scan rate, 0.5 and 0.1 s/frame, respectively. (*Middle*) HS-AFM images of SSP_GD_. For the top and bottom images, operational parameters were: scanning range, 150 × 150 nm (100 × 100 pixels) and 80 × 64 nm (80 × 80 pixels); scan rate, 0.2 and 0.07 s/frame, respectively. (*Right*) HS-AFM images of SSP_GD222_. For the top and bottom images, operational parameters were: scanning range, 200 × 200 nm (100 × 100 pixels) and 80 × 65 nm (80 × 65 pixels); scan rate, 1 and 0.07 s/frame, respectively.

### The seesaw balance can be altered by buffer conditions and ligand binding

Finally, we aimed to alter the seesaw balance of SSP_GD_ using factors other than mutations. We found buffer conditions that could reversibly shift the seesaw balance of SSP_GD_ (Fig. 5A). Through limited proteolysis and DHFR activity measurements, we found that lower salt concentrations and high pH level favored an exclusive GFP state in SSP_GD_, resulting in the loss of DHFR activity (Fig. 5B). Conversely, exchanging the buffer to lower pH and increase the salt concentrations significantly enhanced DHFR activity while reducing GFP fluorescence (Fig. 5B). This shift in SSP_GD_ balance due to buffer exchange was observed repeatedly (Fig. 5B). Additionally, HS-AFM images confirmed that SSP_GD_ exhibited distinct two-domain shapes under DHFR-biased conditions (pH 6 and 500 mM NaCl) (Fig. 5C, Movies S5, S6).

**Figure 5.**
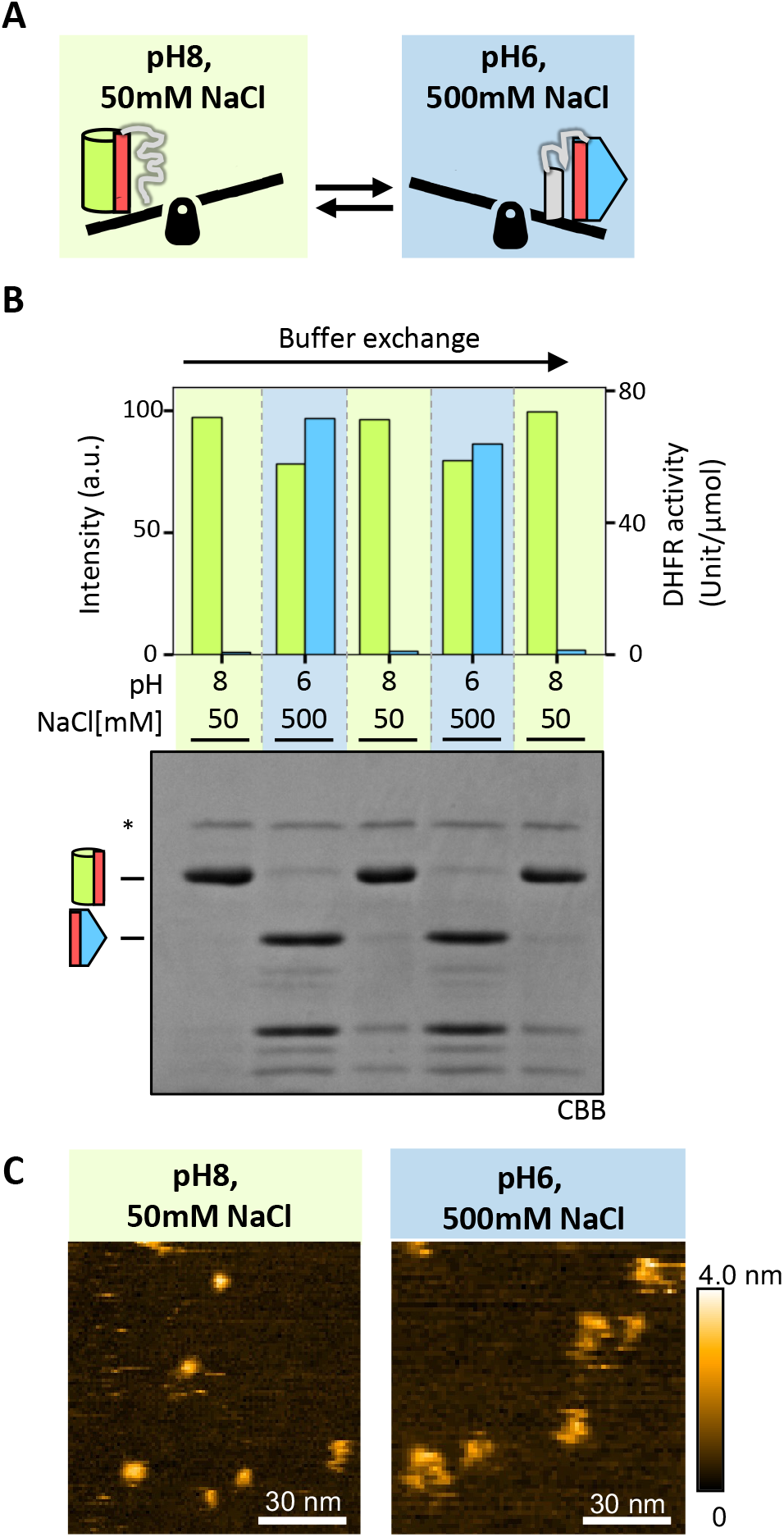
Reversible alteration of seesaw balance by buffer conditions. (*A*) Transition between two SSP_GD_ states by adjusting pH and NaCl concentration. High pH and low NaCl concentration stabilize the GFP-biased state, while low pH and high NaCl concentration stabilize the DHFR-biased state. (*B*) Alteration of SSP_GD_ conformation through buffer exchange. SSP_GD_ was diluted to 100 µM using either a GFP-stabilized buffer (25 mM NaPi pH 8, 50 mM NaCl) or a DHFR-stabilized buffer (25 mM NaPi pH 6, 500 mM NaCl). The buffer conditions were alternated through the cycle of concentration and dilution. Light-colored backgrounds represent the respective conditions (green for GFP-biased and blue for DHFR-biased). GFP fluorescence (*green*) and the DHFR enzymatic activity (*blue*) were measured. The conformational states of SSP_GD_ under each condition were verified by limited proteolysis, as shown in Fig. 3*C*. The asterisk (*) indicates the position of proteinase K. (*C*) HS-AFM imaging of SSP_GD_ structural dynamics under the two buffer conditions. *Left*, HS-AFM imaging of the SSP_GD_ in 25 mM NaPi (pH8.0), 50 mM NaCl. Operational parameters: scanning range, 100 × 100 nm (100 × 100 pixels); scan rate, 0.34 s/frame. *Right*, HS-AFM imaging of the SSP in 25 mM NaPi (pH 6.0), 500 mM NaCl. Operational parameters: scanning range, 150 × 150 nm (100 × 100 pixels); scan rate, 0.2 s/frame.

Next, we added MTX to probe the native state of the DHFR moiety of SSP_GD_. Evaluation of the GFP-to-DHFR state ratio in the presence of MTX through limited proteolysis showed a gradual shift from 9:1 to approximately 1:9 after 4 h (*SI Appendix*, Fig. S3A, B). As MTX specifically binds to the native DHFR state (34, 35), which is nearly irreversible within the timeframe employed here, the kinetics likely reflect the transition between the two states in a dynamic equilibrium.

## Discussion

Here, we proposed the concept of a fusion protein that adopts two interconvertible functional conformations, termed seesaw protein (SSP). This was specifically engineered by fusing GFP, which lacks the C-terminal β-strand, and DHFR, which lacks the N-terminal β-strand, with an overlapping linker (SSP_GD_). Various in vivo to in vitro analyses, including HS-AFM visualization, support the presence of two distinct states in SSP_GD._

Previous studies have shown that proteins with drastic morphological switches exist in nature or can be artificially created by de novo design (7, 10–14, 23–25). SSP_GD_ can be considered an addition to the lineage of these metamorphic proteins, with the following distinguishing features: First, two distinct proteins with completely different functions and topologies were fused; second, their different functions were designed to be evaluated in vivo; and finally, a peptide of only 13 amino acids as an overlapping region (OR) in SSP_GD_ was sufficient to tune the seesaw balance.

Proteins that switch conformations have been designed by dividing the structures of GFP (32, 38, 39) and ribonuclease barnase (31) into two regions, duplicating one region and fusing it to the end, and then replacing the unduplicated region. However, there has been no attempt to design a protein with completely different conformations that compete for the shared region.

The key factors that enabled SSP_GD_ to exhibit seesaw-like behavior include the choice of proteins and the optimization of the OR. GFP and DHFR, both extensively studied, are monomeric, functional proteins with established systems for measuring function in vivo (40). Both proteins are well expressed in *E. coli* and fold efficiently. They are robust to mutagenesis and truncation; for example, Ala insertion at the N-terminus of DHFR has little effect on folding efficiency (41), indicating mutation tolerance when DHFR is placed on the C-terminal side in SSP. GFP is suitable for SSP because the split GFP system allows the C-terminal β-strand (β11) and the other region (β1-10) to be expressed separately and then assembled. Additionally, GFP loses fluorescence if truncated at 226 residues and does not emit fluorescence if the OR is not incorporated (42).

Another key point is the optimization of the OR. In this design, the OR was primarily DHFR-like in sequence to allow SSP to have two different functions: the DHFR activity within SSP_GD_ was almost identical to that of the corresponding monomer, whereas the fluorescent properties of GFP were considerably compromised. For example, it has been reported that the E222Q mutant of eGFP slows the maturation of fluorescence by 1/25 and stabilizes the three-dimensional structure (43, 44). This may explain why GFP fluorescence in SSP_GD_ is only ∼2% of that of wild-type sfGFP.

This impaired GFP moiety is also linked to another issue. When examining the reversible seesaw balance change under altered buffer conditions (Fig. 5B), the entire protein structures were switched as evaluated by limited proteolysis, but the decrease in GFP fluorescence was only about 20%. This suggests that the GFP moiety of SSP_GD_ comprises two populations: a minor fluorescent GFP moiety, which is relatively resistant to seesaw changes, and a major non-fluorescent GFP moiety, which changes the balance easily.

In the future, the SSP principle could be employed to create protein switches within single polypeptide chains that exclusively toggle between completely different functions.

## Materials and Methods

### Plasmid construction and cloning

Plasmids were constructed by inserting genes into the corresponding sites of the pET21c(+) vector using standard cloning procedures and Gibson assembly. Each plasmid was amplified in *E. coli* in LB medium containing appropriate antibiotics. Wild-type GFP lacking 229-238 in the C-terminus, which does not decrease fluorescence intensity or change excitation and emission peak (42), was used for SSP_GD_ and its variants. Superfolder GFP, evolved from wild-type GFP and lacking residues 230-238 in the C-terminus, was used as full-length GFP in this study. An *E. coli* DHFR mutant, ASDHFR (DHFR-C85A/C152S), functionally and structurally equivalent to wild-type DHFR (45), was used to engineer SSP_GD_ and its variants. The SSP mutants, SSP_GD218_ and SSP_GD222_, were engineered by site-directed mutagenesis. Fusion proteins, where full-length GFP and DHFR were fused with a Gly-Ser-Gly-Gly-Gly-Gly-Ser linker, were used for control experiments. For SSP purification, proteins were tagged with 6×His at the C-termini.

### in vivo analysis

*E. coli* BL21(DE3) cells transformed with plasmids were cultured on LB agar plates. Five colonies were selected and tranferred to LB medium, then incubated at 37 °C for 6 h. The cultures were streaked on LB agar plates containing 0.1 mM IPTG and 0, 1, 4 µg/ml TMP, then incubated at 18 °C for 72 h. Fluorescence at 510 nm of the colonies on plates was detected using a LAS4000 image analyzer (Fujifilm).

To assess protein expression and solubility in vivo, cells grown to an OD_600_ of approximately 0.5 at 37 °C were further cultured overnight at 18 °C with 0.1 mM IPTG. Harvested cells were suspended in lysis buffer (20 mM Tris-HCl pH 8.0 and 1 mM EDTA). Total lysates, obtained by sonication (Branson), were centrifuged at 15,000 g for 5 min. The supernatants and the uncentrifuged lysates were subjected to SDS-PAGE. The gel was stained with CBB and imaged using a LAS4000 image analyzer (Fujifilm).

### Protein purification

To purify the sfGFP, DHFR, SSP_GD_, SSP_GD218_, and SSP_GD222_, these proteins were overexpressed in *E. coli* BL21(DE3) cells under the same conditions as described above. The harvested cells were lysed by sonication in 20 mM imidazole buffer (50 mM Tris-HCl pH 7.5, 20 mM imidazole, 200 mM NaCl, and 1 mM DTT). Since all proteins were soluble in *E. coli* cells, supernatants of the lysates were collected by low-speed centrifugation (10,000 g, 5 min, 4 °C) to remove the pellet fraction. Subsequently, another centrifugation at high speed (30,000 rpm, 30 min, 4 °C) was performed to eliminate the remaining pellet. The supernatants were then applied to Ni-NTA agarose (Qiagen) columns that had been pre-equilibrated with 20 mM imidazole buffer. The column was washed with 10 column volumes of 20 mM imidazole buffer. Proteins were eluted using a step gradient of 40-200 mM imidazole in 50 mM Tris-HCl pH 7.5, 200 mM NaCl, and 1 mM DTT. The eluted fractions were analyzed by SDS-PAGE. The buffer content of the fractions containing the target proteins was exchanged to HKM buffer (25 mM HEPES-KOH pH 7.5, 100 mM KCl, 5 mM MgCl_2_) through repeated concentration and dilution using an ultrafiltration apparatus. The purified proteins were then stored at -80 °C. Protein concentrations were determined by absorbance at 280 nm using Jasco V-750 spectrophotometer, with estimated extinction coefficients obtained from Expasy ProtParam.

### Measurement of GFP fluorescence and DHFR enzyme activity

GFP fluorescence was measured as previously described (33). Purified proteins were diluted in HKMD buffer (25 mM HEPES-KOH pH7.5, 100 mM KCl, 5 mM MgCl_2_, 1 mM DTT) to a final concentration of 0.25 µM in a 96-well plate. Fluorescence intensity at 513 nm was measured with an excitation wavelength of 480 nm. DHFR activity was measured as previously described (46). Purified proteins were added to a 96-well plate at a final concentration of 0.2 µM in HKMD buffer containing 400 µM dihydrofolate (DHF) and 400 µM NADPH. The concentrations of DHF and NADPH were determined by absorbance at 282 nm and 340 nm, respectively. The decrease in NADPH was measured spectroscopically at 340 nm (ε340 = 11,800 M^-1^ cm^-1^). Both fluorescence and enzyme activity measurements were performed using a Varioskan LUX Multimode Microplate Reader (Thermo Fisher Scientific). Following these activity measurements, absorbance at 280 nm (protein concentration) and 490 nm (GFP chromophore) was measured on a Jasco V-750 spectrophotometer. Dilution errors were corrected using the estimated molar absorption coefficient.

### Limited proteolysis

Purified proteins were diluted to 5 µM in HKMD buffer. Proteinase K (Merck) was added to a final concentration of 1 µg/ml and incubated at 18 °C for 40 min. The reaction was then quenched with phenylmethylsulfonyl fluoride (PMSF) at a final concentration of 5 mM and separated by SDS-PAGE. The gels were stained with CBB. For N-terminal amino acid sequencing of proteolyzed protein fragments, proteins were transferred onto a polyvinylidene difluoride membrane and analyzed with a gas phase peptide sequencer (PPSQ-21, Shimadzu).

To monitor the transition of SSP_GD_ to a DHFR-biased state, SSP_GD_ (10 µM) was incubated in HKMD buffer in the presence of 50 µM MTX at 30 °C. At the indicated times, aliquots were transferred to 18 °C for 1 min, and proteinase K was then added to a final concentration of 10 µg/ml for 5 min. PMSF was added to a final concentration of 5 mM to quench proteolysis, and the proteins were separated by SDS-PAGE. Gels were stained with CBB and photographed using a LAS4000 image analyzer (Fujifilm).

### Biolayer interferometry (BLI)

The binding of proteins to LaG16 or MTX was measured using the Octet K2 (Pall FortéBio, Menlo Park, CA). All measurements were conducted at 30 °C with an agitation rate of 1,000 rpm. An Amine Reactive Second-Generation (AR2G) sensor was used to measure the interaction between LaG16 and SSPs. LaG16 were immobilized onto the AR2G biosensor as follows: each AR2G sensor was equilibrated in NaOAC pH 6 buffer for 2 min, then activated with a buffer containing 20 mM 1-ethyl-3-(3-dimethylaminopropyl)carbodiimide hydrochloride (EDC) and 10 mM N-hydroxysuccinimide. The activated sensors were modified with the amine group of 1 µM LaG16, then quenched with 500 mM Tris-HCl pH 7.5. After establishing a baseline in Kinetics buffer (PBS supplemented with 0.1% BSA and 0.02% Tween20), the sensor was dipped into Kinetics buffer containing 50 nM SSPs for 600 s, then into Kinetics buffer without proteins for 300 s. A streptavidin (SA) sensor was used for the protein-MTX binding experiment. The sensor was constructed by loading biotin-NH_2_ onto the SA sensor, followed by a reaction in 10 mM NaOAC (pH 6) containing 1 mM MTX and 20 mM EDC, leading to the conjugation of the carboxy group of MTX to the amino group derived from biotin-NH_2_ or SA. The sensor was dipped into Kinetics buffer containing 2 µM GFPΔC_OR, SSP_GD_, SSP_GD222_, or 20 nM SSP_GD218_ for 300 s, then into Kinetics buffer without proteins for 300 s.

### HS-AFM observation of SSPs

AFM imaging of the SSPs in solution was performed using a laboratory-built HS-AFM setup (47, 48). Immobilization of SSPs for AFM imaging was achieved via a His-tag introduced at the C-terminus of the SSPs and a Ni-coated mica. Two microliters of 100 nM SSP_GD_ or the mutants were placed on a Ni-coated mica substrate and incubated for 10 min at room temperature (22−28 °C). Unattached molecules were removed by washing with observation buffer (HKM buffer or 100 mM sodium phosphate (NaPi) pH 6.5, 500 mM NaCl). Imaging was carried out in tapping mode using small cantilevers (BL-AC10DS-A2 or custom-made BL-AC7DS-KU4; Olympus, Tokyo, Japan). The cantilever-free oscillation amplitude was approximately 1.5 nm, and the set-point amplitude was 80−90% of the free oscillation amplitude. The imaging rate, scan size, and feedback parameters were optimized to enable visualization using a minimum tip force.

### Buffer exchange experiment

SSP_GD_ was diluted to 100 µM with a GFP-stabilized buffer (25 mM NaPi pH 8.0, 50 mM NaCl) or a DHFR-stabilized buffer (25 mM NaPi pH 6.0, 500 mM NaCl). The buffer containing SSP_GD_ was exchanged with a different-state-stabilized buffer on a NAP-5 column (Cytiva). SSP_GD_ was then concentrated by ultrafiltration. Buffer exchange and protein concentration were repeated alternately. SSP_GD_ concentrations were measured by absorbance at 280 nm. For the limited proteolysis experiment, SSP_GD_ was diluted to 5 µM with the buffer. Proteinase K was added a final concentration of 10 µg/ml and incubated for 3 min at room temperature. The reaction was then quenched with PMSF (5 mM final concentration), separated by SDS-PAGE, and stained with CBB. GFP fluorescence and DHFR enzyme activity of the proteolyzed products were measured as described above, but with a high buffering capacity buffer (400 mM HEPES-KOH pH 7.5, 100 mM KCl, 5 mM MgCl_2_) to maintain constant pH.

### Statistical Analysis

Student’s *t*-test was used to calculate statistical significance, with a two-tailed distribution and unequal variance. All experiments were conducted at least three times independently, and mean values ± standard deviation (SD) are represented in the figures.

## Acknowledgments

We thank Keisuke Yoshida and Toru Hisabori for peptide sequencing; Zhu Bo for BLI; Toshio Ando (Kanazawa University) for providing us with HS-AFM instruments; Integrative Biosciences and the Biomaterials Analysis Division, Open Facility Center at the Tokyo Institute of Technology for DNA sequencing. This work was supported by MEXT Grants-in-Aid for Scientific Research (Grant Numbers JP18H03984, JP20H05925, and JP21H04763 to HT), the World Premier International Research Center Initiative (WPI), MEXT, Japan, and the Kanazawa University CHOZEN project to HK.

