## supporting figures for "Seesaw protein: Design of a protein that adopts interconvertible alternative functional conformations"

**This PDF file includes:**

Supplementary Figures S1 to S3

Legends to Supplementary Movies S1 to S5

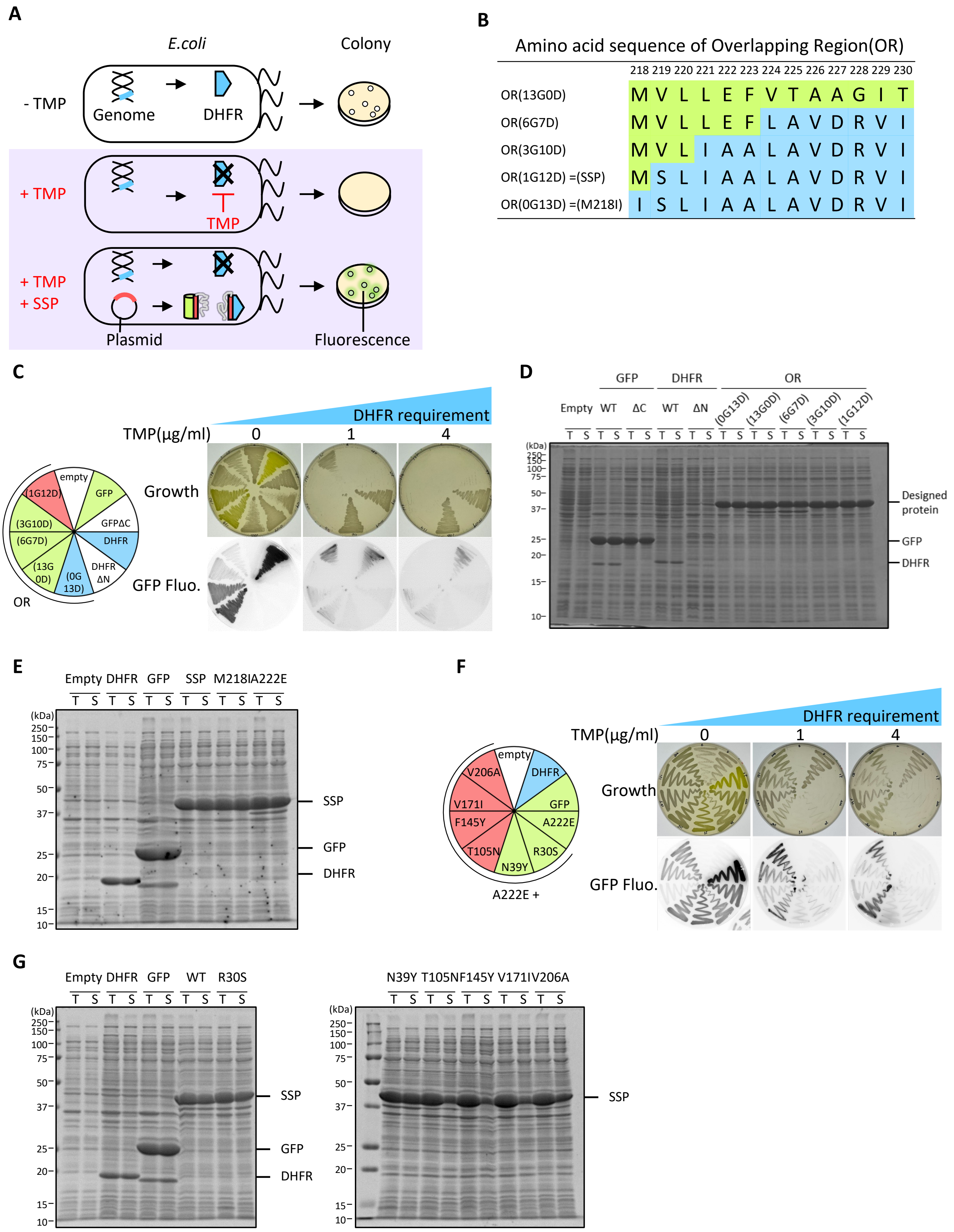
A
E.coli
Colony
- TMP
Genome
DHFR
+ TMP
TMP
+ TMP
+ SSP
Plasmid
Fluorescence
B
Amino acid sequence of Overlapping Region(OR)
218 219 220 221 222 223 224 225 226 227 228 229 230
OR(13G0D) M V L L E F V T A A G I T
OR(6G7D) M V L L E F L A V D R V I
OR(3G10D) M V L I A A L A V D R V I
OR(1G12D) =(SSP) M S L I A A L A V D R V I
OR(0G13D) =(M218I) I S L I A A L A V D R V I
C
DHFR requirement
TMP(μg/ml) 0 1 4
Growth
GFP Fluo.
empty
(1G12D)
GFP
(3G10D)
GFPΔC
(6G7D)
DHFR
(13G 0D)
(0G 13D)
DHFR ΔN
OR
D
GFP DHFR OR
Empty WT ΔC WT ΔN (0G13D) (13G0D) (6G7D) (3G10D) (1G12D)
T S T S T S T S T S T S T S T S T S T S
(kDa)
250 150 100 75 50 37 25 20 15 10
Designed protein
GFP
DHFR
E
Empty DHFR GFP SSP M218IA222E
T S T S T S T S T S T S
(kDa)
250 150 100 75 50 37 25 20 15 10
SSP
GFP
DHFR
F
DHFR requirement
TMP(μg/ml) 0 1 4
Growth
GFP Fluo.
empty
DHFR
V206A
V171I
GFP
A222E
F145Y
T105N
R30S
N39Y
A222E +
G
Empty DHFR GFP WT R30S
T S T S T S T S T S
(kDa)
250 150 100 75 50 37 25 20 15 10
SSP
GFP
DHFR
N39Y T105N F145Y V171I V206A
T S T S T S T S T S
(kDa)
250 150 100 75 50 37 25 20 15 10
SSP

**Fig. S1. Design of SSPs with two functional conformations.**

(***A***) Principle of colony assay: *E. coli* requires functional DHFR for cell division and growth (*top*). *E. coli* with endogenous DHFR cannot grow on plates containing trimethoprim (TMP), a DHFR inhibitor (*middle*). Functional DHFR from SSPs complements the growth defect, enabling *E. coli* growth (bottom). Correct folding of the GFP moiety in SSPs results in fluorescence within the colony (*bottom*). (***B***) Amino acid sequences of the overlapping region (OR) for SSP design. Amino acids from GFP and DHFR are depicted in green and blue, respectively. Numbers and letters in brackets indicate the number of GFP- and DHFR-derived amino acids in the OR sequence. For example, (6G7D) denotes six N-terminal amino acids from GFP and seven amino acids from DHFR. (***C***) Colony assay to assess SSP functionality. The position of overexpressed proteins in cells is depicted in the circle (*left*). "*Empty*" refers to cells without protein overexpression. Proteins were overexpressed for 72 h at 18 °C on plates with varying TMP concentrations (0, 1, and 4 μg/ml). Plates were photographed (*top*) and imaged for fluorescence (*bottom*). (***D***), (***E***) Expression and solubility of SSP candidates (*D*) and the SSP mutants (*E*). *E. coli* lysates expressing SSP candidates were centrifuged, and supernatant fractions were electrophoresed and stained with CBB. 'T' and 'S' denote total and supernatant fractions, respectively. Although most proteins were efficiently expressed and soluble, DHFR_ΔN was not expressed in *E. coli.* (***F***) Colony assay to assess GFP-destabilizing mutations (R30S, N39Y, T105N, F145Y, V171I, V206A) in $SSP_{GD222}$. (***G***) Expression and solubility analysis of SSP mutants from (*F*).

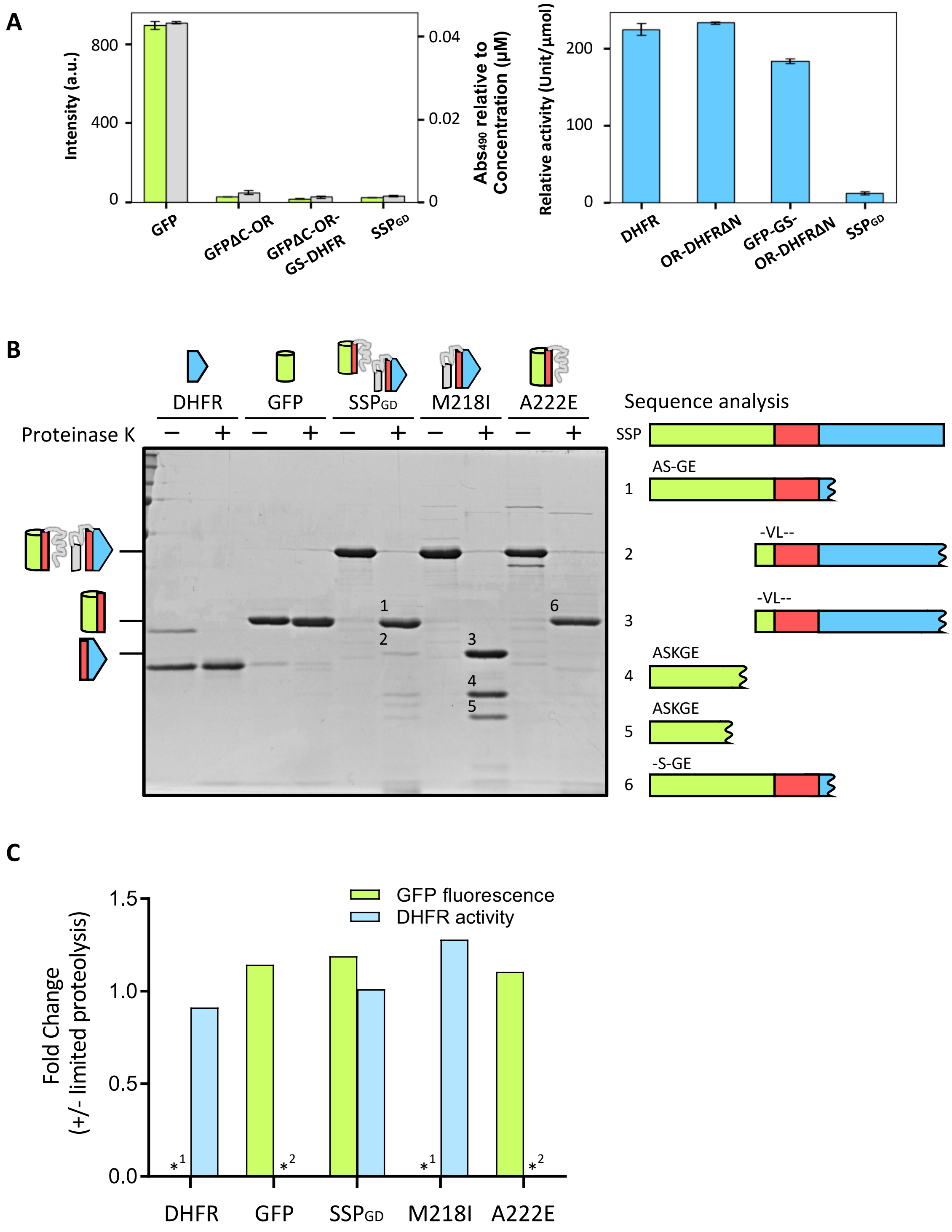


**Fig. S2. Biochemical analysis of purified SSPs.**

(***A***) Evaluation of DHFR enzymatic activity (*left*) and GFP fluorescence (*right*) of SSP$_{GD}$, its parent monomers (DHFR and GFP), and the derivatives. "*GS*" denotes the linker composed of Gly-Ser-Gly-Gly-Gly-Gly-Ser between the GFP and DHFR regions. All measurements were performed at room temperature. For GFP fluorescence, excitation and emission wavelength were 480 and 513 nm, respectively (*green*, *left axis*). The absorption of chromophore was measured at 490 nm (*gray, right axis*). Unit is defined as the amount of enzyme degrading 1 μmol of substrate per minute at room temperature. Data represent the means (±S.D.) of three independent experiments. (***B***) N-terminal sequencing of proteolyzed products, as shown in Fig. 3*C*. The N-terminal sequences of the proteins numbered in the gel image were analyzed using Edman degradation, depicted on the right. Estimated proteolyzed products are illustrated in the schematic of GFP-ΔC (*green*), OR (*red*), and ΔN-DHFR (*blue*). (***C***) Assessment of GFP fluorescence and DHFR enzyme activity after proteinase K treatment. Changes in fluorescence intensity and enzyme activity were compared with and without protease treatment. “*1” and “*2” indicate cases where GFP fluorescence or DHFR activity was below the detection limit.

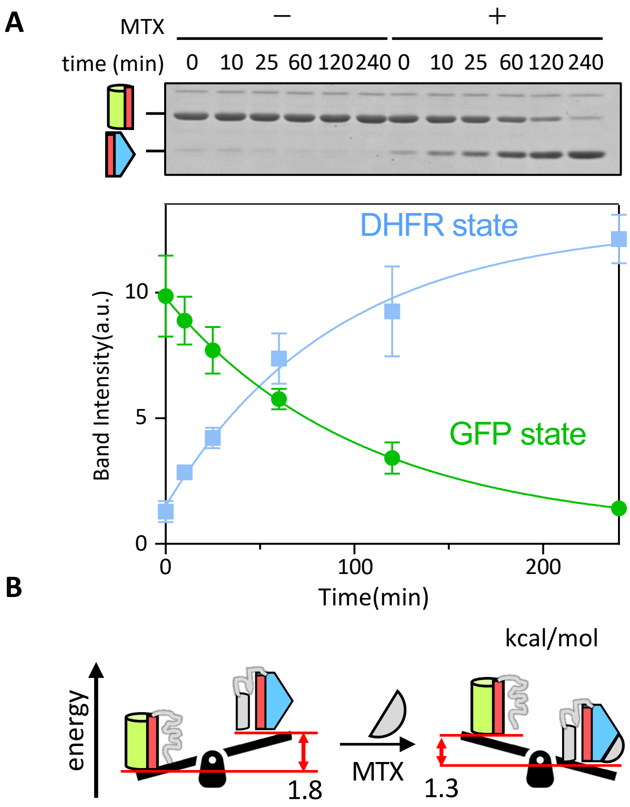


**Fig. S3. Transition of $SSP_{GD}$ to the DHFR-biased state investigated by MTX binding.** (***A***) $SSP_{GD}$ was incubated with (+) or without (-) MTX for specified durations at 30 °C and then subjected to proteinase K treatment. The conformations of $SSP_{GD}$ were monitored through SDS-PAGE and CBB staining (*top*). The band intensities of the two states in the presence of MTX were quantified and plotted (*bottom*). Solid lines represent the change in band intensity fitted by single exponential kinetics. Data represent the means (±S.D.) of three independent experiments. (***B***) Alteration in the seesaw slope induced by MTX. The free energy difference between the two states of $SSP_{GD}$ was determined from the band intensity of limited proteolysis, as in Fig. 3D. In the presence of MTX, this difference was calculated assuming equilibrium after 4 h of incubation and that all DHFR-states bound to MTX.

**Legends to Supplementary Movies**

**Movie S1. HS-AFM movie showing the structural dynamics of $SSP_{GD222}$.**

The operational parameters were scanning range, 80 × 65 nm (80 × 65 pixels); scan rate, 0.07 s/frame.

**Movie S2. HS-AFM movie showing the structural dynamics of $SSP_{GD218}$.**

Scanning range, 70 × 700 nm (70 × 70 pixels); scan rate, 0.1 s/frame.

**Movie S3. HS-AFM movie showing the structural dynamics of $SSP_{GD}$.**

The operational parameters were scanning range, 80 × 64 nm (80 × 80 pixels); scan rate, 0.07 s/frame.

**Movie S4. HS-AFM movie showing the multiple $SSP_{GD}$ at a wide range**

Scanning range, 150 × 150 nm (100 × 100 pixels); scan rate, 0.2 s/frame.

**Movie S5. HS-AFM movie showing the structural state of $SSP_{GD}$ in a GFP-biased buffer (pH 8.0, 50 mM NaCl).**

Scanning range, 100 × 100 nm (100 × 100 pixels); scan rate, 0.34 s/frame.

**Movie S6. HS-AFM movie showing the structural state of $SSP_{GD}$ in a DHFR-biased buffer (pH 6.0, 500 mM NaCl).**

Scanning range, 150 × 150 nm (100 × 100 pixels); scan rate, 0.2 s/frame.
